# Regionally specific and highly pleiotropic loci together mediate skeletal evolution in tropical and temperate house mice

**DOI:** 10.1101/2025.10.03.680311

**Authors:** Sylvia M. Durkin, Kathleen G. Ferris, Mallory A. Ballinger, Ke Bi, Gabriela Heyer, Michael W. Nachman

## Abstract

When organisms are exposed to new environments, they often evolve a suite of adaptive phenotypic changes. Understanding the genetic basis of these changes gives us insight into the factors that constrain and facilitate the process of adaptation. Here we use quantitative trait locus (QTL) mapping in wild-derived house mice from temperate and tropical environments to describe the genetic architecture of adaptive skeletal evolution. This study provides a unique view into the evolution of discordant phenotypic changes within the interconnected skeletal system, since tropical and temperate mice have evolved under the opposing forces of Allen’s and Bergmann’s rules (shorter extremities, but larger body size in cold climates). We generated an F3 mapping population of 449 mice and measured a variety of cranial and post-cranial skeletal traits. First, we discovered that temperate and tropical house mice have undergone extensive skeletal divergence in accordance with Allen’s and Bergmann’s rules. Warm-adapted mice have longer limbs, longer pelvic girdles, longer and narrower skulls, but smaller overall body size. Second, we identified 83 QTL across 13 traits and found that the genetic basis of skeletal divergence involved mainly additive, small-effect loci distributed across the genome. While most QTL influenced only one or two traits, we also identified several highly pleiotropic QTL that influenced multiple traits across the skeletal system, often antagonistically to the evolved difference. Such pleiotropic loci may constrain adaptation. Lastly, we found that QTL for overall skeletal variation were enriched for signatures of selection in wild house mouse populations, providing evidence that skeletal variation in this system is indeed driven by adaptive evolution.

**Article Summary:** The incredible diversity of the vertebrate skeleton offers an opportunity to explore the genetic architecture of phenotypic variation, which is essential to understanding the process of evolution. We characterize the genetic basis of morphological evolution using QTL mapping in house mice adapted to differing climates. Skeletal divergence is widespread in these mice and reflects temperature adaptation. This divergence is mediated by small-effect loci scattered across the genome, as well as several highly-pleiotropic loci. Interestingly, pleiotropic loci often influence phenotypes in the opposite direction from the evolved change, perhaps constraining the process of adaptation. This work demonstrates how complex morphological changes can evolve within an interconnected system, such as the skeleton, under functional, developmental, and selective constraints.

## Introduction

Environmental adaptation is a major driver of phenotypic diversity. Describing the proximate and ultimate causes of such variation is a fundamental goal in evolutionary biology. Two widespread ecogeographic patterns that reflect environmental adaptation are Allen’s and Bergmann’s rules, the observations that animals in colder regions have shorter extremities, such as limbs, ears, and tails (Allen 1877), but larger overall body size (Bergmann 1847). This variation represents adaptation to different thermal environments, since animals with large bodies and small extremities have a smaller surface area to volume ratio thereby helping to conserve heat (Mayr 1956). When present together, Allen’s and Bergmann’s rules exemplify oppositional selective forces on morphology since some parts of the body are selected to be larger while other parts of the body are selected to be smaller. Studying populations that have recently adapted to differing climates thus offers a unique means to explore how complex, discordant phenotypic changes evolve at the genetic level and understand how genetic architecture operates to both constrain and facilitate the adaptive process.

Pleiotropy, when a single locus influences more than one trait, is classically thought of as being a constraining force on adaptation, as pleiotropic effects are expected to be largely deleterious (Otto 2004; Fisher 1930). However, pleiotropy may also be synergistic, conferring beneficial changes in more than one trait. In such instances, pleiotropy may facilitate adaptation, especially if populations are further from their fitness optima and must evolve multiple phenotypic changes (Orr 1998; Hämälä et al. 2020). Studies in stickleback fish (Rennison and Peichel 2022), *Arabidopsis* (Frachon et al. 2017), *Mimulus* (Lowry and Willis 2010; Ferris et al. 2017), and ragweed (Hämälä et al. 2020) have found support for intermediate levels of pleiotropy driving adaptive change, but QTL studies mapping large numbers of traits often recover a majority of low-pleiotropy loci, with a smaller number of loci influencing multiple traits (Wagner et al. 2008; Kenney-Hunt et al. 2008; Albert et al. 2008).

On the other hand, modularity in genetic architecture, where single traits or suites of related traits are controlled independently from other evolving phenotypes, may allow organisms to escape the deleterious effects of pleiotropy (Wagner, Pavlicev, and Cheverud 2007). Modular changes can occur through context-dependent mechanisms such as *cis-*regulatory elements that achieve temporal or regional specificity (Stern 2000; Wagner, Pavlicev, and Cheverud 2007; Carroll 2008). Examples ranging from the developmental specificity of the *Hox* gene family across vertebrates (Carroll 2005; 2008) to regionally specific control of stickleback skeletal traits (Miller et al. 2014) suggest that modularity may be a common feature in trait evolution.

House mice (*Mus musculus domesticus*) in the Americas provide a tractable model to study the genetic basis of environmental adaptation as they have expanded into a broad range of habitats across North and South America from their ancestral range in Western Europe within the past 500 years (Phifer-Rixey and Nachman 2015; Agwamba and Nachman 2023). In this short time, mice in different regions have adapted to different environments through changes in morphology, physiology, and behavior (Phifer-Rixey et al. 2018; Ballinger et al. 2023; Dumont et al. 2024), including variation in overall body size and extremity length, with populations from northern latitudes being larger (Bergmann’s rule) and having relatively shorter limbs, ears, and tails (Allen’s rule) (Lynch 1992; Phifer-Rixey et al. 2018; Ferris et al. 2021; Ballinger and Nachman 2022). Here we use variation in skeletal traits to explore the genetic architecture of these adaptive phenotypes. Importantly, the interconnected nature of the skeleton allows us to ask how multiple phenotypic changes evolve simultaneously within a system under both functional and developmental constraints (Cheverud 1996; Lande 1979; Schluter 1996; Klingenberg 2008). Considerable work has been done in connecting genotype to phenotype for naturally occurring skeletal differences in contexts such as house mouse body size (Parmenter et al. 2022; Payseur et al. 2023; Parmenter et al. 2016; Wilches et al. 2021), deer mouse tail length variation (Hager et al. 2022; Kingsley et al. 2024), craniofacial differences in African cichlids (Powder and Albertson 2016), predator defense phenotypes in stickleback (Albert et al. 2008; Miller et al. 2014), and variation in human skeletal form (Kun et al. 2024; Xu et al. 2025).

The contrasting nature of Allen’s and Bergmann’s rules allow us to study pleiotropy and modularity in the context of evolved oppositional changes in the skeleton: mice in colder regions have evolved to be larger but also to have shorter limbs. If individual genes cause all skeletal elements, including limb bones, to be larger, the pleiotropic nature of such genes might constrain adaptation. On the other hand, if different genes generally influence different bones, such a modular genetic architecture could facilitate adaptation. Skeletal development is known to be mediated by both modular and pleiotropic regulatory networks. For example, while T-box transcription factors 4 (*Tbx4*) and 5 (*Tbx5*) both target fibroblast growth factor 10 (*Fgf10*) during bone formation, *Tbx4* is specific to hindlimbs and *Tbx5* is specific to forelimbs (Gibson-Brown et al. 1996; Agarwal et al. 2003; Naiche and Papaioannou 2003; Minguillon et al. 2012). On the other hand, master regulators like runt-related transcription factor 2 (*Runx2*) are crucial for overall skeletal development and are expressed widely in skeletal and non-skeletal tissue (Jeong et al. 2008; Takarada et al. 2016; Komori 2018). The study of skeletal traits also makes it possible to address foundational issues in quantitative genetics concerning the distribution of effect sizes and dominance patterns expected during adaptation.

In this article, we use QTL mapping to understand the genetic architecture of skeletal divergence between two groups of locally adapted house mice from Manaus, Brazil (tropical) and Saratoga Springs, New York (temperate). First, we found that temperate and tropical mice show divergence across all regions of the skeleton, in accordance with Allen’s and Bergmann’s rules. Skeletal divergence is mediated by many small-effect, additive loci across the genome. However, there are multiple highly pleiotropic QTL that influence many traits across the skeleton. These QTL frequently act in opposition to the evolved phenotypic difference, perhaps constraining the process of adaptation. Lastly, we show that skeletal QTL are enriched for signatures of natural selection in wild populations of house mice. Together these results speak to how complex adaptive changes are achieved within the interconnected parts of the skeleton.

## Materials and Methods

### Ethics statement

This work was conducted in accordance with the University of California, Berkeley Institutional Animal Care and Use Committee (AUP-2017-08-10248 and AUP-2016-03-8548). Euthanasia was performed under approval of the relevant ACUC at the University of California, Berkeley, and included the humane use of isoflurane and cervical dislocation by trained personnel.

### Animal husbandry and mice used

All mice were housed under a 12:12 light cycle at 70-72°F and fed standard rodent food *ablibitum*. Wild derived, partially-inbred strains of house mice (*Mus musculus domesticus*) used in this study were originally collected from 43°N in Saratoga Springs, New York (SARA) and at 3°S in Manaus, Brazil (MANA). Information on the establishment of these inbred strains is described in Phifer-Rixey et al. (2018) and Ferris et al. (2021). In brief, live mice were collected from Saratoga Springs and Manaus, brought back to the lab, and unrelated individuals were mated to create the N1 generation. Inbred strains were generated via subsequent sib-sib mating. SARA and MANA mice used in this study were inbred for four generations, which is expected to reduce the heterozygosity of the mouse strains by 60% (Silver 1995).

### QTL intercrossing experiment

To generate a QTL mapping panel, MANA and SARA mice were crossed in both directions (i.e MANA females X SARA males, and vice versa) to produce the F1 generation. F1 mice were intercrossed to produce 100 F2 mice, which were then intercrossed to produce an F3 mapping panel of 449 mice with mixed SARA and MANA genotypes.

### Phenotyping

At 10 weeks of age (69-72 days), F3 mice were euthanized and body weight measurements were taken. Additionally, liver tissue was collected in >95% pure ethanol from all F3 mice and stored at -80°C for later DNA extraction and sequencing. Skeletons were prepared from all individuals as standard museum specimens and deposited in the mammal collection of the UC Berkeley Museum of Vertebrate Zoology (catalog numbers are listed in Supplementary File 1). Skeletal phenotypes were measured on cleaned specimens (Figure 1B). Appendicular measurements including humerus length, ulna length, femur length, tibia length, pelvic length, pelvic width, and metatarsal length were taken using digital calipers (Mitutoyo Digimatic Caliper #500-150-30). Measurements for each appendicular element were taken three times and the mean of the three measures was used for analysis. Cranial measurements including skull length, skull width, bizygomatic width, zygomatic length, rostrum length, rostrum width at the premaxilla, and maximum rostrum width were collected with a Keyence digital microscope (VHX-970FN), specifically using the point-to-point measurement tool.

**Figure 1.**
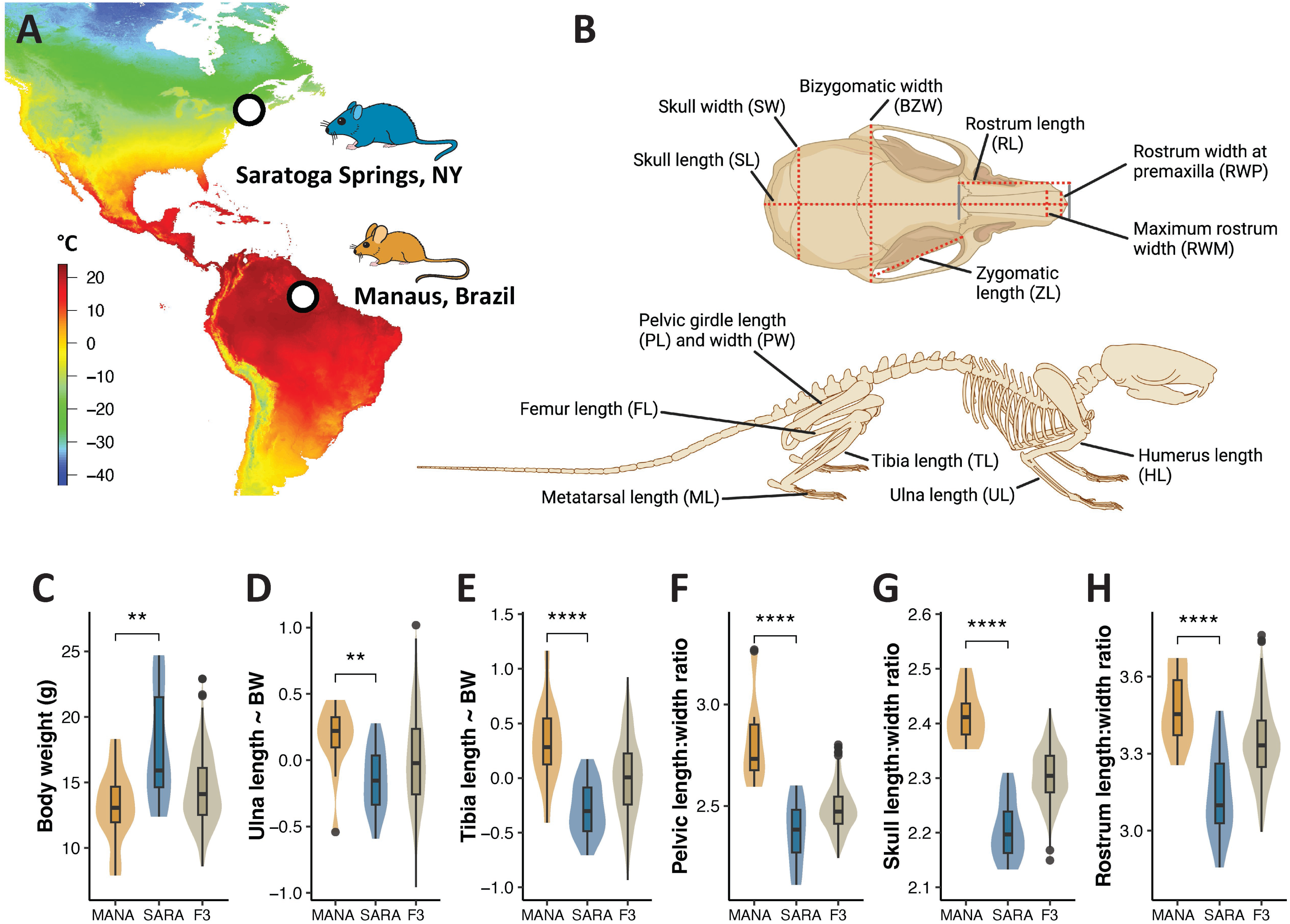
Extensive skeletal divergence in MANA and SARA mice in accordance with Allen’s and Bergmann’s rules. A). Minimum annual temperature at original sampling localities for SARA and MANA inbred lines. Panel adapted from Durkin et al. (2024). B). Names and acronyms for all skeletal measurements collected. C-H). Phenotypic variation in MANA, SARA, and F3 hybrid mice. Panels D and E represent residual values based on the correlation of the focal phenotype and body weight. Significance between MANA and SARA calculated with Welch two sample t-tests; * = p < 0.05, ** = p < 0.01, *** = p < 0.001, **** = p < 0.0001.

A total of 14 skeletal measurements were taken; details on landmarks used for each measurement are in Supplementary File 2. For parental phenotypes, we measured SARA and MANA mice from the same generation of inbreeding as the parents of the QTL cross (MANA = 14 mice, 6 females, 8 males; SARA = 16 mice, 7 females, 9 males). For parental mice, we additionally calculated multiple allometry metrics including the intermembral index (ratio of the forelimb to hindlimb [(HL + UL)/(FL + TL) x 100]), humerofemoral index (ratio of proximal elements of the limb [(HL/FL) x 100]), crural index (ratio of distal to proximal elements of the hindlimb [(TL/FL) x 100]), and brachial index (ratio of distal to proximal elements of the forelimb [(UL/HL) x 100]). Divergence between MANA and SARA and subsequent p-values were determined using Welch two sample t-tests.

### Sequencing and Genotyping

We genotyped parent and F3 mice using double-digest restriction site-associated (ddRAD) sequencing (Peterson et al. 2012). First, we collected liver tissue at 10 weeks of age from all 449 F3 mice and 14 SARA and MANA parents. Genomic DNA was extracted using Qiagen DNeasy Blood and Tissue kit, and digested with MSP1 and EcoR1. End-specific P1 and P2 adapters were ligated and library preparation was undertaken as described in Peterson et al. (2012). Samples were pooled in equimolar ratios and sequenced across three lanes of Illumina HiSeq4000. Two samples had duplicate names at this stage, and were removed from downstream analysis, bringing the total number of F3 individuals to 447.

Reads were aligned to the *Mus musculus* reference genome (GRCm39) using BWA mem (bwa version 0.7, Li and Durbin 2009). We called variants for all F3 and parental mice using the Genome Analysis Toolkit (GATK version 4.1.0, Depristo et al. 2011). HaplotypeCaller was run on all individuals separately. Resulting gVCFs were merged using CombineGVCFs, and then genotyped together using GenotypeGVCFs. To define a set of marker SNPs for QTL mapping, we filtered for quality by depth (QD > 5), genotype quality (GQ > 20), and minor allele frequency (MAF > 5%) using vcftools (vcftools version 0.1.14, Danecek et al. 2011). We then removed variants that were identical across F3 mice using GATK SelectVariants and missing in more than 60% of individuals using vcftools. Then, we filtered for segregation distortion by removing autosomal SNPs not in Hardy-Weinberg equilibrium among the F3 mice using vcftools (--hwe 0.05). Lastly, we selected markers that had fixed and different genotypes between the SARA and MANA parent individuals. Through this filtering, we identified 3,521 high quality markers to be used for QTL mapping.

Genotypes for the F3 mice were converted to ABH format using TASSEL 5 (Bradbury et al. 2007), which denotes whether the individual is homozygous for SARA allele (A), homozygous for the MANA allele (B) or heterozygous (H). We estimated an initial genetic map from the physical distances of our 3,521 markers using mmconvert in R, which is based on the Cox et al. mouse genetic map (Cox et al. 2009).

### Genetic map construction

Prior to QTL mapping, we further filtered markers and individuals for genotype missingness and redundancy. Specifically, we removed individuals with fewer than 100 genotyped markers using the subset function in R/qtl (Broman and Sen 2009), which retained 437 F3 mice. We then removed redundant markers using findDupMarkers and drop.markers, resulting in a total of 2,778 markers. We then re-estimated the genetic map originally inferred using the Cox et al. (2009) map by calculating intermarker distances using the Carter-Falconer map function in est.map in R/qtl with a genotyping error rate of 0.2%. Lastly, we visually inspected the resulting genetic map using plotMap in R/qtl and removed problematic markers that skewed the genetic map due to incorrect recombination estimates using droponemarker, which resulted in a final 2,777 markers for the final downstream analysis.

### QTL Identification

We identified traits that showed significant differences between SARA and MANA mice and then identified regions of the genome associated with variation in these traits (Supplementary File 3, zygomatic length and metatarsal length were not mapped, as they were not different between SARA and MANA mice) using R/qtl (Broman and Sen 2009). We performed both single QTL, and multi-QTL mapping. For single QTL mapping, we performed a genome scan for association at 1cM step points using the scanone function in R/qtl, specifying the Haley-Knott method. For skeletal traits, we included sex, body weight, and collector ID as additive covariates, and for body weight, only sex was included as an additive covariate. We performed permutation tests separately for each trait in R/qtl using the scanone function (n = 1000 permutations on the autosomes and n = 27,560 permutations on the X chromosome), and QTL were considered significant at the 5% genome-wide significance threshold.

Multiple QTL mapping was performed using the scantwo function in R/qtl, specifying the Haley-Knott method. This approach identifies additional QTL that often do not rise to genome-wide significance in single QTL mapping by searching for two QTL simultaneously, controlling for the phenotypic effect of one and then scanning for additional QTL (Broman and Sen 2009). Analogous covariates were used as in single QTL mapping. We used the scantwo LOD significance thresholds determined for a mouse intercross specified in Broman and Sen (2009) calculated from 10,000 simulations of crosses with 250 individuals, markers at a 10 cM spacing, and analysis by Haley-Knott regression (full model = 9.1, conditional-interactive model = 7.1, interaction model = 6.3, additive model = 6.3, and conditional-additive model = 3.3). QTL in the scantwo search were considered significant if the test for the additive QTL model and the test for the association between the second identified QTL and phenotype being mapped was significant (conditional-additive model). Importantly, all QTL identified with the single-QTL scanone method were also identified with the multi-QTL scantwo method.

To determine the number of skeletal traits influenced by a single QTL, we defined overlapping QTL using the upper and lower interval of the peak (1.5 LOD on either side of the peak) implemented with lodint in R/qtl. Overlapping QTL were designated with the same QTL ID (listed in Supplementary File 4), thereby facilitating analyses of QTL that influence multiple phenotypes. Importantly, QTL with non-overlapping intervals were never classified with the same QTL ID, even if there was a larger QTL that spanned both non-overlapping QTL, which was common for chromosome 13 (Figure 2). In these instances, the larger QTL spanning the two smaller QTL was classified into the QTL ID of whichever smaller QTL spanned the LOD peak for the larger QTL.

**Figure 2.**
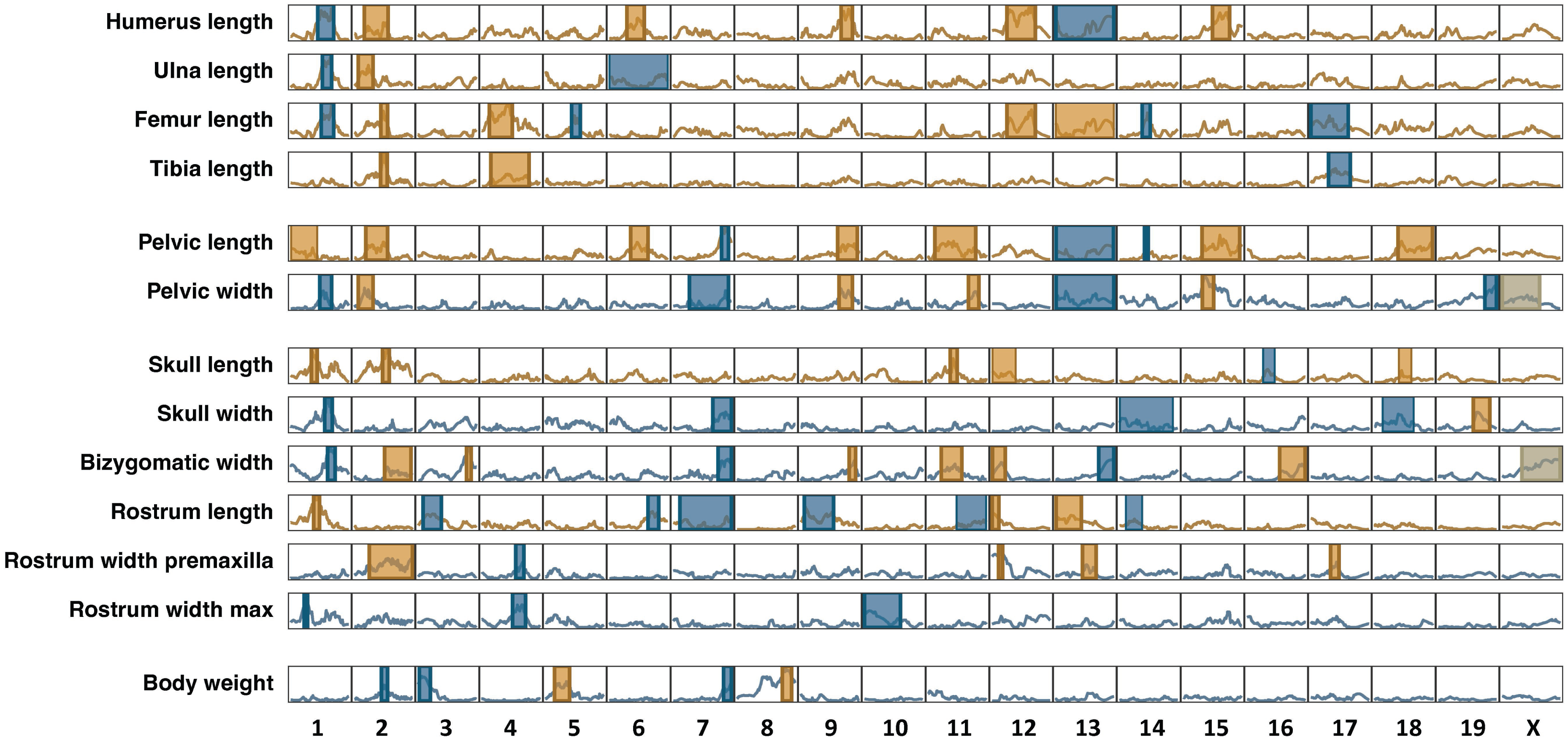
QTL for skeletal divergence in SARA and MANA mice. Shaded regions represent the confidence window around each QTL peak (1.5 LOD on either side of the peak). QTL intervals are shaded according to whether the SARA allele (blue) or MANA allele (yellow) increases the phenotypic value. The background LOD score line for each trait corresponds to whether SARA (blue) or MANA (yellow) has the larger phenotypic value. QTL effects cannot be estimated on the X chromosome in R/qtl and are shaded grey.

### Genetic effects

We measured genetic effects of skeletal and body weight QTL, including additivity, dominance, effect size, and percent variance explained with R/qtl. We first constructed a multi-QTL object for each trait containing all significant QTL using makeqtl. We then used fitqtl to calculate dominance effects (d), additive effects (a), and percent variance explained for each QTL. Additive effects, also referred to as effect size, represents half the phenotypic difference between F3 mice homozygous for the MANA allele and F3 mice homozygous for the SARA allele. Dominance effects represent the difference between the phenotypic mean of heterozygous F3 mice and the phenotypic mean of SARA and MANA homozygous F3 mice together. Effect sizes (a) were standardized by dividing by the standard deviation of the individual traits (a/sd), and dominance effects (d) were then standardized by dividing by a (d/a). We performed a goodness of fit test to assess whether the distribution of effect sizes fit an exponential curve using the LK.test function in the EWGoF R package.

We then categorized QTL into seven categories based on the value of the d/a ratio as defined by Kenney-Hunt et al. (2006): strong underdominant (d/a < -2.5), underdominant (-2.5 < d/a < -1.5), SARA dominant (-1.5 < d/a < -0.5), additive (-0.5 > d/a < 0.5), MANA dominant (1.5 > d/a > 0.5), overdominant (2.5 > d/a > 1.5), and strong overdominant (d/a > 2.5).

### Clustering of highly pleiotropic QTL (hpQTL)

To determine if overlapping QTL mapped to more phenotypes than expected by chance, we used a custom python script (available on GitHub: <u>s-durkin/Durkin_skeletalQTL_2025</u>) that reshuffles QTL windows to test the expected degree of overlap among identified QTL. More specifically, for each trait, the identified QTL windows were shuffled on the genome, while accounting for the length and gene density of individual chromosomes (ie. longer and more gene rich chromosomes are more likely to have QTL). After all trait QTL were re-placed on the genome, we calculated the number of pairwise overlaps between QTL windows. This was done 1,000 times to simulate the expected number of QTL overlaps per chromosome. We calculated a p-value of significant QTL clustering by taking the proportion of the simulated pairwise overlap distribution that was outside the actual number of pairwise overlaps for each chromosome. We performed this test separately for QTL identified through single and multi-QTL mapping approaches. QTL on chromosomes that had more clustering than expected by chance are referred to as highly pleiotropic QTL (hpQTL).

We next evaluated the influence of recombination rate on QTL identification and effect size. We estimated recombination rates for the QTL cross in 5Mb sliding windows using the est.recrate function in the R/xoi package, which estimates recombination rate across a chromosome (at 0.25cM intervals) using the physical and genetic map of a QTL cross. Details about the physical and genetic map for our study are in the *Sequencing and Genotyping* and *Genetic Map Construction* sections above. We then used the estimated recombination rates to 1) compare the average recombination rate within all QTLs on a given chromosome to the average recombination rate of that chromosome, and 2) compare the average recombination rate within a QTL to the effect size of that QTL. To evaluate if the recombination rate of QTLs on a given chromosome were higher than the chromosome average, we calculated a p-value for each chromosome representing the proportion of the distribution of estimated recombination rates (at each 0.25cM step) that was beyond the average recombination rate within the QTLs on that chromosome. We evaluated the relationship between the average recombination rate within a QTL and its absolute effect size using a Pearson correlation.

### Principal Component Analysis and Mapping

We performed a principal component analysis (PCA) for parents and F3 mice together on all skeletal traits (except zygomatic length and metatarsal length) and body weight. Briefly, we used the prcomp function in R/stats and removed F3 mice that did not have complete data for all skeletal measurements. We then performed single-QTL and multi-QTL mapping for PC1 and PC2 analogously as described above. Because we wanted to include the maximum number of individuals for QTL mapping, we imputed PC values for F3 mice that had missing skeletal data using the pca function in R/pcaMethods, specifying the “ppca” method, and extracted imputed PC values for all F3 individuals using the completeObs function in R/pcaMethods (Stacklies et al. 2007). Missing data was the result of damaged bones during the skeleton preparation process, and constituted 0.7% of all measurements. We ran scanone and scantwo analyses twice each for PC1 and PC2, once with body weight, sex, and collector ID as additive covariates, and once with just sex and collector ID as additive covariates.

### Phenotypic and QTL sharing correlations

We calculated phenotypic correlations among all F3 mice for all skeletal traits (except zygomatic length and metatarsal length) and body weight using the cor function in the R/stats package, specifying the Pearson correlation method. We then connected the phenotypic correlations to the degree of QTL sharing in two ways. First, we constructed a matrix of QTL sharing, recorded as a proportion of QTL shared, for each pairwise trait comparison using the QTL IDs defined as stated in the *QTL Identification* section above. We then calculated a Pearson’s correlation between the proportion of QTL shared between two traits and the phenotypic correlation between those two traits. Second, we calculated a Pearson’s correlation between the average phenotypic correlation for a trait across all its pairwise trait correlations, and the average proportion of QTL shared for a trait across all pairwise comparisons.

### PBSn1 tests for selection and overlap with QTL peaks

To connect the QTL mapping results to signatures of selection in wild populations of house mice, we used previously published results of a normalized version of the population branch statistic (PBSn1) from Durkin et al. (2024). The PBS test identifies allele frequency shifts that are present in one focal population as compared to two outgroups, and are therefore candidates for loci under selection (Yi et al. 2010; Crawford et al. 2017). We used two published PBSn1 tests, one where a population from Manaus, Brazil is the focal group, and one where a population from the border between New Hampshire and Vermont (NH/VT) is the focal group. The NH/VT population is geographically close to Saratoga Springs, the original sampling locality of the SARA line. In both tests, Iran and France were the outgroup populations, which are near the ancestral range of *Mus musculus domesticus* (Morgan et al. 2022; Phifer-Rixey and Nachman 2015). These tests identified genes containing SNPs that were in the top 1%, 5%, and 10% of the distribution of PBSn1 scores, which were used for downstream analysis.

To assess whether genes within skeletal QTL peaks were enriched for signatures of selection, we looked at the overlap between genes in the top 1%, 5%, and 10% of PBSn1 scores and genes under QTL peaks for PC1 and PC2. We chose to look at PC peaks as they capture the overall genetic basis of skeletal divergence between SARA and MANA mice. We tested whether this overlap was more than expected by chance by simulating the expected overlap between randomly chosen gene sets the same size of the PBSn1 gene set and the PC peaks gene set. We then calculated the overlap between 10,000 iterations of randomly sampled gene sets and tabulated a p-value using the proportion of the simulated distribution that was outside the observed overlap value. This was done on PBSn1 outliers that were present in either the Manaus or NH/VT test at each outlier cutoff level (1%, 5%, and 10%), for a total of three overlap enrichment tests.

## Results

### Temperate and tropical mice show morphological divergence across the skeleton in accordance with Allen’s and Bergmann’s rules

As reported previously, MANA mice are smaller than SARA mice by an average of 4.4 g, representing 33% of MANA’s body weight (Figure 1C, Supplementary File 3) (Ballinger and Nachman 2022; Durkin, Ballinger, and Nachman 2024; Ballinger et al. 2023). MANA are also known to have longer ears and tails (Ballinger and Nachman 2022). To assess whether tropical and temperate house mice differ in skeletal traits in accordance with Allen’s and Bergmann’s rules, we measured 14 phenotypes in SARA and MANA mice across skull, pelvis, and limb bones (Figure 1 A-B, Supplementary Files 1-2). To assess divergence between SARA and MANA, we focused on 1) differences between the raw phenotype values and 2) differences between the residuals of each measurement’s correlation with body weight to control for the variation in skeletal traits that is simply a function of body size. We did not use relative measures as these are often reflective of body weight differences themselves, as opposed to correcting for differences in body weight (Packard and Boardman 1988).

We found that MANA mice have longer limb bones including the humerus, ulna, femur, and tibia and have a longer pelvic girdle as compared to SARA mice (p < 0.05 for residuals, Figure 1C-F, Supplementary File 3). MANA mice also have a relatively longer and narrower skull and rostrum (p < 0.05 for residuals, Figure 1G-H, Supplementary File 3). Additionally, we assessed difference in skeletal shape using allometry indices including the intermembral index (ratio of the forelimb to hindlimb), humerofemoral index (ratio of proximal elements of the limb), crural index (ratio of distal to proximal elements of the hindlimb), and brachial index (ratio of distal to proximal elements of the forelimb). We found that MANA mice show a disproportionate elongation of distal limb elements (p < 0.05 for crural and brachial indices, Supplementary Figure 1). Distal elements of the skeleton have been hypothesized to be more evolutionarily labile (Rothier et al. 2023), and are more exposed to the external environment, resulting in greater responses of distal skeletal elements to temperature variation (Serrat 2014). Overall, tropical and temperate mice showed substantial divergence in skeletal phenotypes across the cranial and axial skeleton that are reflective of adaptation to different temperatures consistent with Allen’s and Bergmann’s rules.

### Skeletal phenotypes are mediated by many additive, small effect QTL across the genome

To understand the genetic basis of skeletal divergence in tropical and temperate mice, we performed a QTL intercrossing experiment between SARA and MANA resulting in a mapping panel of 449 F3 mice. High-quality genetic markers used for QTL mapping (2,777 total) had an average intermarker distance of 0.86cM and average depth of 28.4 reads. Skeletal phenotypes for these F3 hybrid mice were intermediate between SARA and MANA distributions (Figure 1C-H). We mapped body weight and 12 skeletal phenotypes (zygomatic length and metatarsal length were removed as they did not show divergence between SARA and MANA; Supplementary File 3) using a single-QTL and multi-QTL mapping approach. Unless otherwise noted, we report results from multi-QTL methods; single-QTL mapping results are reported in the supplement.

We identified 83 QTL across the 12 skeletal traits and body weight that were distributed across every chromosome in the genome. The number of QTL identified per trait ranged from three to ten, and the total variance explained per trait, summed across all QTL for that trait, was between 4.7% to 12.8% (Figure 2, Supplementary File 4). Individual QTL explained a small percent of phenotypic difference, with the average being 1.7%, the largest explaining 4.7% of phenotypic variation (QTL for maximum rostrum width) and the lowest explaining only 0.3% (QTL for humerus length) (Figure 3A, Supplementary Figure 2). The distribution of percent variance explained for all QTL fits an exponential distribution (Cox-Oakes goodness of fit test, p-value < 0.001), as predicted by Orr (1998). To better understand the influence of individual QTL on phenotype, we calculated standardized additive and dominance effects for each QTL allowing us to estimate effect size and mode of inheritance. Standardized effect sizes for each QTL were small (Figure 3B), influencing skull phenotypes an average of 0.03 mm and axial skeletal phenotypes an average of 0.08 mm (Supplementary File 4). Further, by calculating standardized dominance values (d/a) we found that 51% of QTL are additive (d/a between -0.5 and 0.5, Figure 3C), while 35% of QTL were either SARA allele or MANA allele dominant (d/a between -1.5 and -0.5 and d/a between 0.5 and 1.5, respectively). We also found that additive QTL had larger effect sizes (Figure 3D), a pattern that has been reported in other studies investigating the genetic basis of skeletal evolution (Miller et al. 2014).

**Figure 3.**
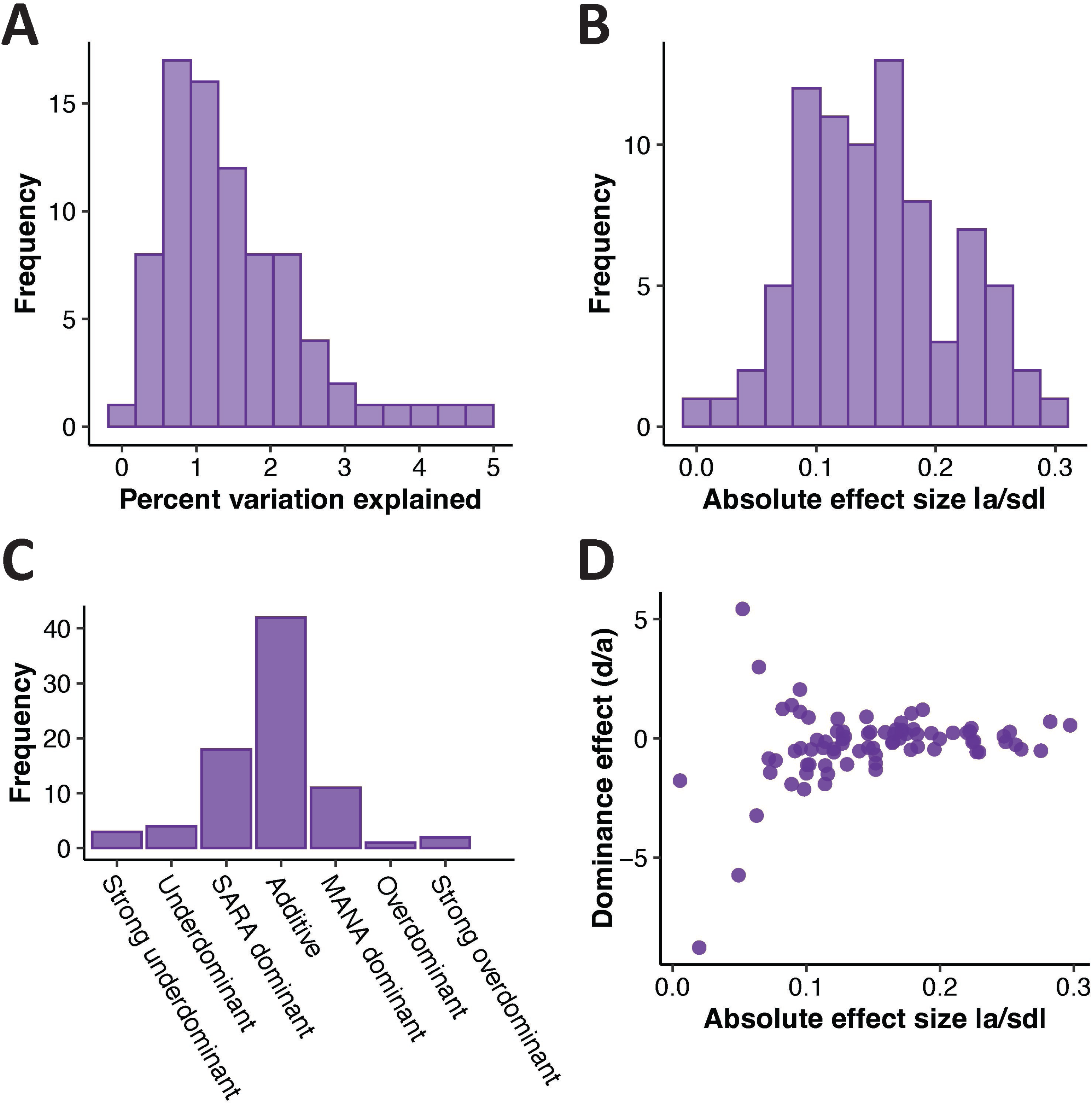
QTL for skeletal variation are mostly additive and of small effect. A). Percent variation explained for all individual QTL. B). Absolute standardized effect size for all individual QTL. C). Dominance patterns across QTL. Dominance categories were defined as follows: strong underdominant (d/a < -2.5), underdominant (-2.5 < d/a > -1.5), SARA dominant (-1.5 > d/a < -0.5), additive (-0.5 > d/a < 0.5), MANA dominant (1.5 < d/a > 0.5), overdominant (2.5 > d/a < 1.5), and strong overdominant (d/a > 2.5). D). Relationship between dominance and effect size for all individual QTL.

Lastly, we classified QTL as being concordant or antagonistic to the direction of evolutionary change (ie. MANA alleles conferring a larger trait value when that trait is larger in MANA would be concordant, but if that trait is larger in SARA mice, it would be antagonistic), and found that QTL often influence phenotypes in the opposite direction of the evolved difference. For example, femur length, which is relatively longer in MANA mice, has four QTL in which the MANA allele makes the trait longer, and four QTL where the SARA allele makes the trait longer (Figure 2). Overall, only 54% of QTL operate in the same direction as the evolved phenotypic difference between SARA and MANA mice.

### The genetic architecture of skeletal divergence is largely regionally specific, but marked by several highly pleiotropic QTL

Overall, we found low levels of pleiotropy with most QTL influencing only one or two traits (Figure 4A). For example, over two-thirds of the QTL affected only 15% of the traits measured, consistent with previous reports of low levels of pleiotropy for skeletal traits in mice (Wagner et al. 2008). At the same time, there were several genomic regions that harbored QTL influencing multiple traits. Specifically, QTL on chromosomes 1, 2, 7, and 9 each influenced four or more separate traits (Figure 4A, details on defining overlapping QTL in *Materials and Methods*). These QTL comprise 27% of all individual QTL identified (22 of 83 total QTL). To evaluate whether these QTL are mapping to more phenotypes than expected by chance, we simulated the expected number of pairwise QTL overlaps per chromosome and compared that to the observed number of pairwise overlaps. We found that three chromosomes, 1, 2 and 13, had more overlapping QTL than expected by chance (Figure 4B). Importantly, when looking at QTL identified through the single-QTL mapping approach only, chromosomes 1 and 2 had an enrichment of QTL overlaps, whereas chromosome 13 did not (Supplementary Figure 3). Further, many QTL on chromosome 13 span the entirety of the chromosome (Figure 2), so QTL on that chromosome may overlap even when the peaks of the QTL are far apart. For these reasons, we focus primarily on the overlapping QTL on chromosomes 1 and 2, referred to as highly pleiotropic QTL (hpQTL).

**Figure 4.**
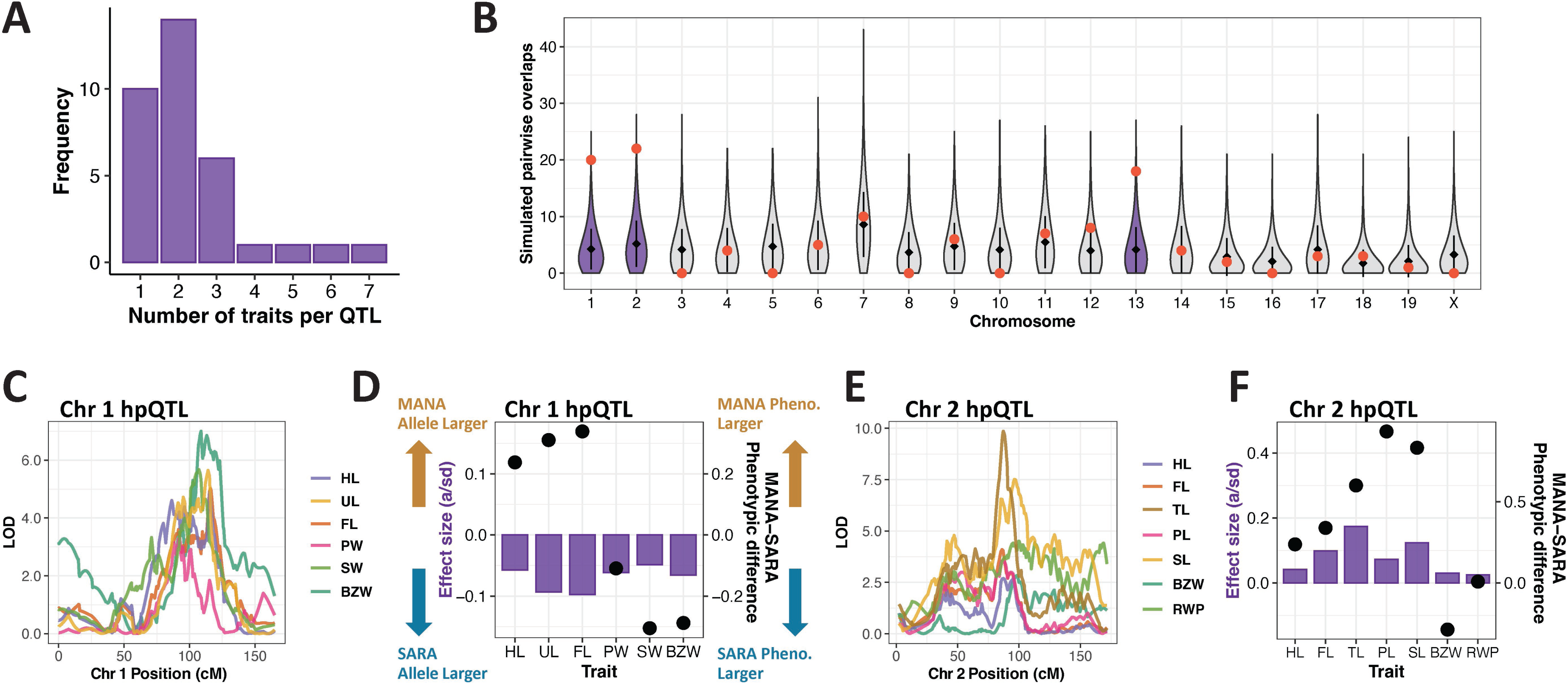
Highly pleiotropic QTL are involved in skeletal evolution. A). Number of traits influenced per QTL. B). Simulated number of pairwise QTL overlaps per chromosome. Red dot represents the observed value of pairwise overlaps. Chromosomes that contained more overlaps than expected by chance (observed value beyond the top 5% of simulated distribution) are colored purple. C). Overlapping LOD curves for the phenotypes influenced by the hpQTL on chromosome 1. D). Mode of action for chromosome 1 hpQTL. Left y-axis and purple bars display the effect size and direction of the chr 1 hpQTL on each trait it influences; negative values correspond to the SARA allele conferring a higher phenotype value, and vice versa for the MANA allele. Right y-axis and black points display the phenotypic difference between MANA and SARA, for residual phenotype values. For example, at the chr 1 hpQTL the SARA allele makes humerus length (HL) longer as shown by the purple bar, but it is actually MANA mice that have longer HL, as shown by the black dot. E-F). Overlapping LOD curves (E) and mode of action (F) for chromosome 2 hpQTL. Direction of action for MANA/SARA alleles and phenotypes analogous to those shown in panel D.

### Highly pleiotropic QTL influence phenotypes unidirectionally, often opposite to the direction of evolutionary divergence

hpQTL on chromosomes 1 and 2 influence traits across the axial and cranial skeleton (Figure 4C, E), suggesting little regional modularity. Skeletal divergence between SARA and MANA represents a system of oppositional changes (i.e. some trait values are higher in SARA, and some trait values are higher in MANA) since selection has presumably acted to increase overall body size and to decrease the length of the extremities in cold-adapted SARA mice (and vice-versa in warm-adapted MANA mice). To explore how hpQTL influence skeletal traits in this context, we compared the sign of the effect size (negative represents the SARA allele conferring the larger phenotype value, and vice versa for MANA) to the phenotypic difference between MANA and SARA. We found that hpQTL always influence different traits in the same direction, and this direction sometimes does not match the direction of evolutionary phenotypic divergence between SARA and MANA (Figure 4D, F). For example, having the SARA allele at the chromosome 1 hpQTL confers larger trait values for all traits affected by this locus. However, only half of the traits influenced by this hpQTL are larger in SARA mice. This mode of influence is consistent with our results for QTL more generally, since about half of all QTL act in the opposite direction of the evolved change (Figure 2).

While the QTL on chromosomes 7 and 9 did not map to more phenotypes than expected by chance, they still influence traits in the same way as the hpQTL on chromosomes 1 and 2, meaning each trait is influenced in the same direction, and this does not always agree with the direction of evolutionary change (Supplementary Figure 4). Overall, we found that skeletal variation in tropical and temperate mice is mediated by a combination of regionally specific, modular genetic changes and highly pleiotropic loci that influence traits across the entire skeletal system.

### Recombination rate and QTL detection

We next investigated the potential influence of variation in recombination rate on QTL detection. QTL can be localized to specific chromosomal segments more easily in regions of high recombination, owing to the greater number of crossovers per physical distance (Capilla-Pérez et al. 2024), but regions of low recombination may lead to a biased detection of strong QTL owing to linkage over longer physical distances (Noor et al. 2001). By estimating recombination rate between genomic markers in the QTL mapping cross, we were able to confirm that the average recombination rate across QTL intervals is not different from the average background rate (Supplementary Figure 5A). Second, we compared the estimated average recombination rate within QTL regions to the effect size of those QTL and found no significant correlation (Supplementary Figure 5B). Together, these results suggest minimal biases influencing the detection and estimation of effect size for skeletal trait QTL.

### Skeletal traits are highly correlated in F3 mice, potentially due to a shared genetic basis

To evaluate the degree of interconnection across the skeleton, we looked at correlations among all traits and assessed whether these correlations were reflected at the genetic level. First, we found that skeletal phenotypes are highly correlated among F3 mice, and most are strongly correlated with body weight (Figure 5A). All pairwise correlations were significant (p < 0.05, Pearson’s correlation), and R^2^ values ranged between 0.40 and 0.91, with an average of 0.68.

**Figure 5.**
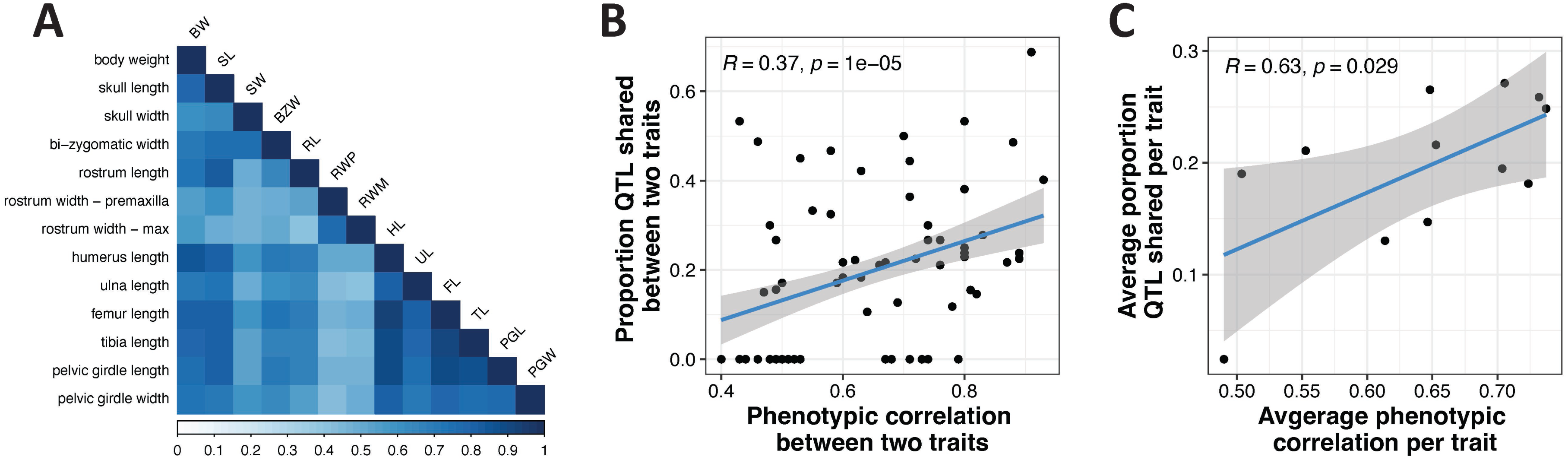
Skeletal phenotypes are highly correlated, in part due to a shared genetic basis. A) Pairwise Pearson’s correlation coefficients for all divergent skeletal traits and body weight. Scale represents Pearson’s R value. B). Relationship between phenotypic correlation and proportion of shared QTL for each pairwise phenotype correlation. C). Same relationship as in panel B, but averaged across all pairwise comparisons for a single trait. Correlation coefficients and p-values for B and C are Pearson’s R.

We next sought to connect the degree of phenotypic correlation to the genetic similarity between traits. To do this we first calculated the proportion of QTL that are shared between each pairwise trait comparison. There is a significant, positive relationship between the phenotypic correlation coefficient between two traits, and the proportion of QTL shared between those traits (Figure 5B). This is also true when looking at the relationship between the phenotypic correlation and proportion of QTL shared when averaged across all pairwise comparisons for a single trait (Figure 5C). Overall, we found that different skeletal traits were highly correlated. Higher phenotypic correlations were associated with increases in QTL sharing, suggesting that the strong correlations among skeletal traits are in part due to a shared genetic basis.

### QTL for principal components map to highly pleiotropic QTL

Because skeletal traits co-vary, we performed a principal components (PC) analysis to reduce the dimensionality of the trait data and to gain insight into how skeletal variation, in aggregate, is reflected at the genetic level. A PC analysis of divergent skeletal measurements for all F3 mice and MANA and SARA separates the two parental groups (Figure 6A). While most F3 mice fall between the parental groups, they do span nearly the full range of variation, including values represented by the parents (Figure 6A). Together, PCs 1 and 2 explain 78.72% of trait variation, with PC1 loading strongly with body weight (Figure 6A, D-E) and PC2 loading with skull phenotypes such as rostrum width and skull width (Figure 6A, F-G). Interestingly, the phenotypes related to PC2 were those with relatively weaker pairwise correlations (Figure 5A). These phenotypes were also less strongly associated with overall body size as compared to other skeletal traits measured (Figure 5A).

**Figure 6.**
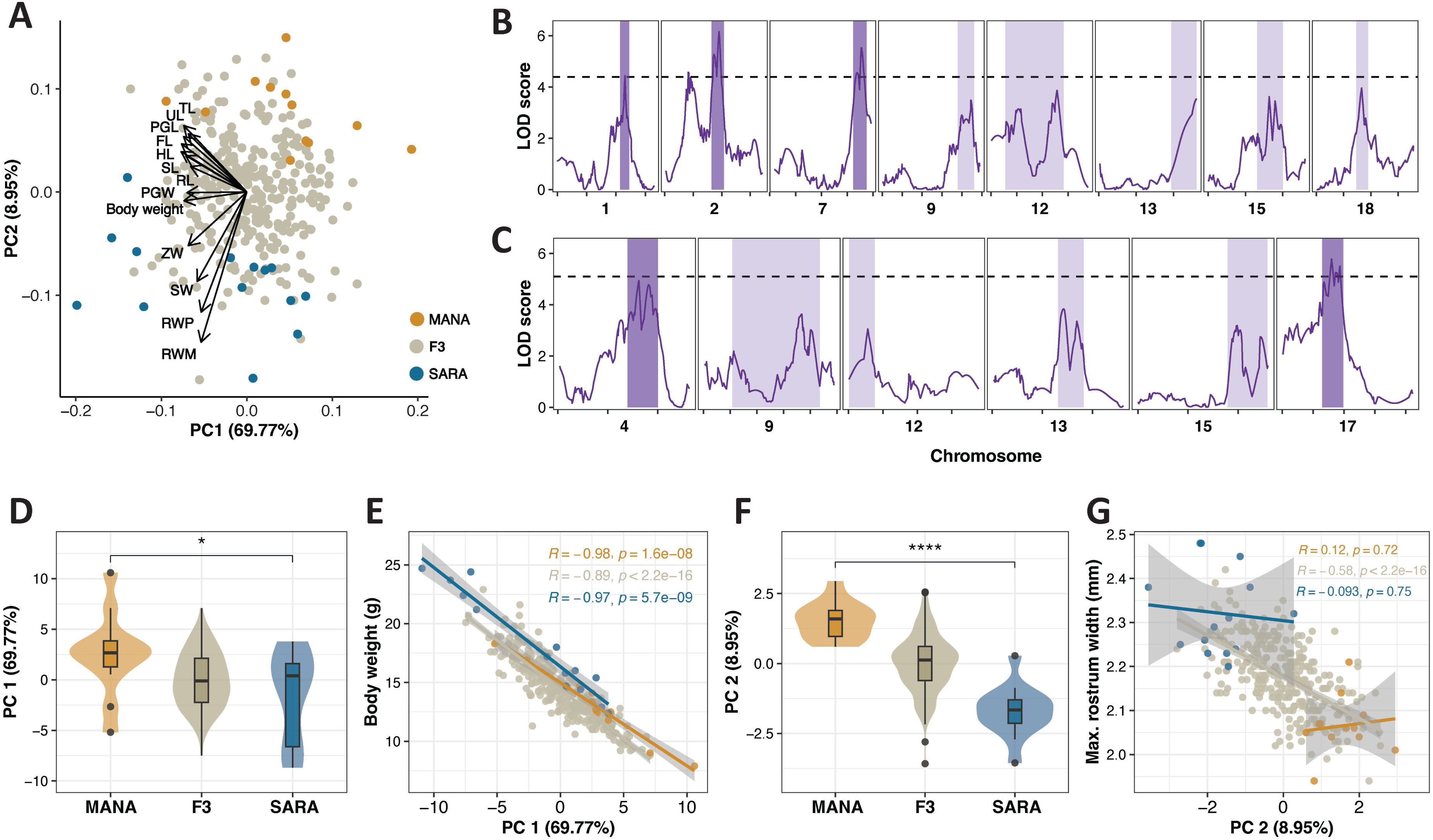
Principal components of skeletal variation are influenced by body size and map to hpQTL. A). PCA of divergent skeletal phenotypes and body weight for MANA, SARA, and F3 hybrid mice. B-C). Significant QTL intervals for PC1 (B) and PC2 (C). Dark purple QTL were identified through the single-QTL mapping approach, while light purple intervals correspond to QTL identified only through the multi-QTL mapping approach. Body weight was included as a covariate for both B and C. D). Distribution of PC1 values for MANA, SARA, and F3 hybrid mice. E). Relationship between PC1 and body weight. Correlation coefficient and p-value are Pearson’s R. F). Distribution of PC2 values for MANA, SARA, and F3 hybrid mice. G). Relationship between PC2 and maximum rostrum width. Correlation coefficient and p-value are Pearson’s R. Significance between MANA and SARA in D and F calculated with Welch two sample t-tests; * = p < 0.05, ** = p < 0.01, *** = p < 0.001, **** = p < 0.0001.

We first identified QTL mapping to PC1 and PC2 using just sex, and not body weight, as a covariate, and found that peaks for PC1 largely overlap QTL for body weight (Supplementary Figure 6A, Figure 2, Supplementary File 4), which agrees with the strong correlation between body weight and PC1. Similarly, peaks for PC2 largely overlap peaks for rostrum width measurements (Supplementary Figure 6B, Figure 2, Supplementary File 4). When body weight is included as a covariate, QTL for PC1 overlap the hpQTL on chromosomes 1 and 2, and the non-significant hpQTL influencing 4 or more traits on chromosomes 7 and 9 (Figure 6B). This reinforces the idea that hpQTL influence many aspects of skeletal divergence. QTL identified for PC2 with body weight included as a covariate were similar to those identified when only sex was used, suggesting that PC2 is not influenced by body weight.

### QTL for principal components are enriched for signatures of selection

To connect our findings on genetic variation in lab-reared animals to natural selection acting on wild populations of house mice, we utilized previously published selection scans from Durkin et al. (2024). Specifically, we used a normalized version of the population branch statistic (PBSn1), which identifies allele frequency shifts that are unique to one focal population in comparison to two outgroup populations. We used two previously published PBSn1 tests, one where Manaus was the focal population, and one where a population on the border of New Hampshire and Vermont (NH/VT) was the focal population. The NH/VT population is geographically close to Saratoga Springs, the original sampling locality of SARA mice. For both tests, France and Iran were outgroup populations, which are near the ancestral range of *Mus musculus domesticus* (Phifer-Rixey and Nachman 2015; Ferris et al. 2021; Morgan et al. 2022). We focused on connecting selection scans to QTL peaks for PC1 and PC2 as these capture a large proportion of phenotypic divergence between MANA and SARA. Moreover, given the large width of some of the QTL windows (Figure 2), testing across all QTL identified would encapsulate much of the entire genome.

We looked for an enrichment of genes in QTL intervals for PC1 and PC2 in the top 5% of PBSn1 outliers. Using permutation tests, we found a greater overlap than expected by chance between PC genes and genes under selection in either NH/VT or Manaus (Figure 7A-B, p-value = 0.005). This enrichment of PC genes among selection scan outliers was significant at the 1%, 5%, and 10% cutoffs of top PBSn1 scores (Supplementary Figure 7). These results are consistent with the idea that skeletal divergence between SARA and MANA mice was in part driven by natural selection acting on the wild populations from which they were derived.

**Figure 7.**
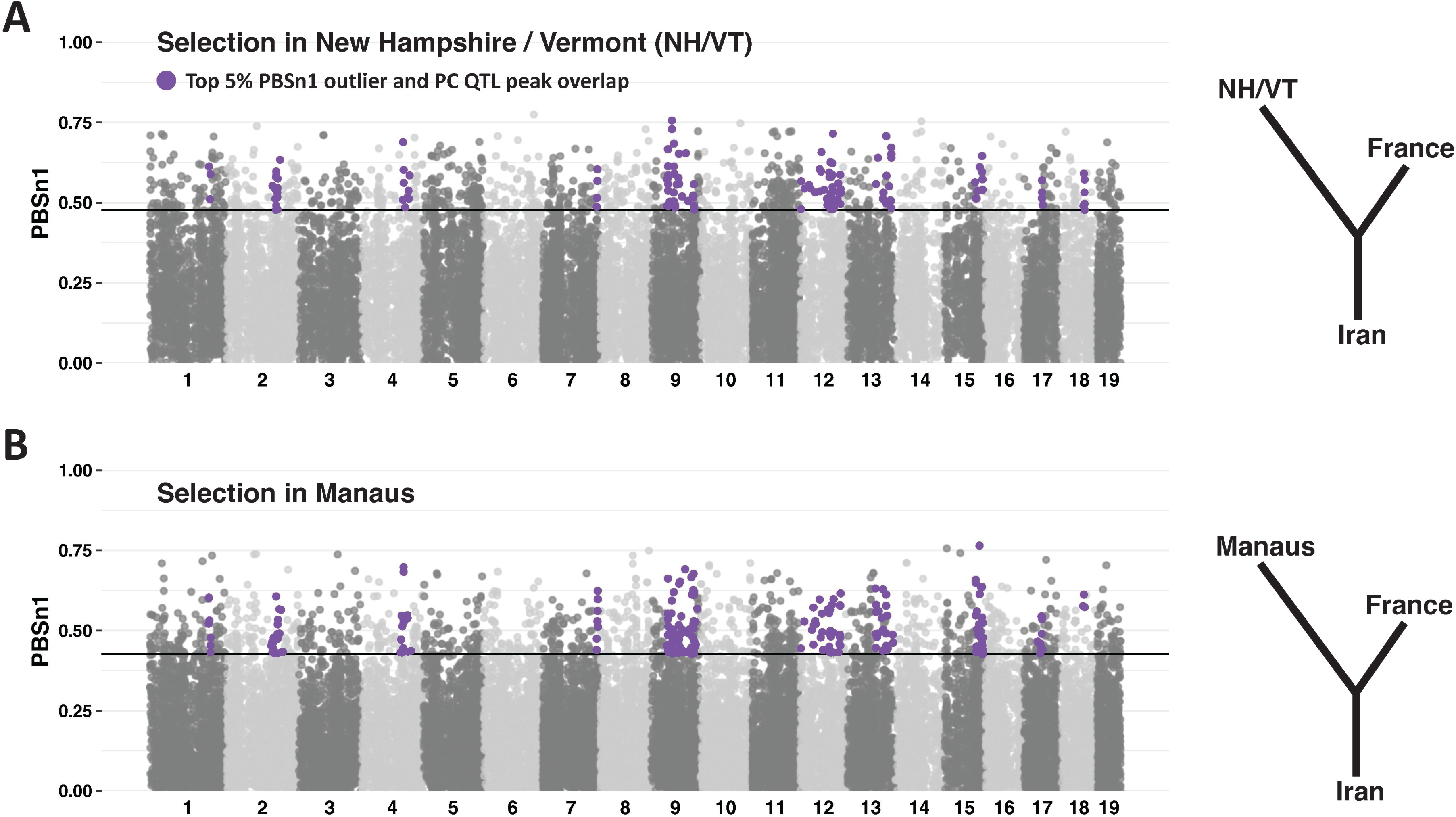
Genes within principal component QTL intervals are enriched for signatures of selection in natural populations. A-B). PBSn1 values for selection scans with New Hampshire/Vermont (NHVT) (A) and Manaus as the focal population (B). France and Iran are included as outgroups in both tests. Line indicates the top 5% outliers. Purple dots are PBSn1 outliers that are within QTL intervals for PCs 1 and 2.

## Discussion

Studying populations that have recently adapted to differing climates allowed us to characterize the genetic basis of adaptive change during the early stages of divergence. We used QTL mapping between tropical and temperate house mice to understand how discordant morphological changes can evolve within the interconnected system of the skeleton. First, we found that SARA and MANA mice show extensive divergence across the entire skeletal system. This divergence is mediated by many small-effect, additive loci scattered across the genome. While most QTL influenced only one to two traits, we did identify several pleiotropic QTL influencing four or more skeletal phenotypes. These pleiotropic QTL map to the same genomic regions as the QTL underlying PCs for skeletal variation which summarize overall phenotypic divergence. Lastly, we found that genes within PC QTL regions were enriched for signatures of selection in wild populations of house mice, suggesting that skeletal differences between SARA and MANA mice were in part shaped by adaptative evolution.

### Adaptation to temperature variation is reflected in the skeletons of temperate and tropical mice

Allen’s (1877) and Bergmann’s (1847) rules are two of the most well-documented ecogeographic patterns in birds (e.g. Johnston and Selander 1964; Nudds and Oswald 2007; Symonds and Tattersall 2010; McQueen et al. 2022) and mammals (e.g. Yom-Tov and Nix 1986; Blackburn, Blackburn, and Hawkins 2004; Alhajeri et al. 2020). Changes in body size and proportions are thought to be thermoregulatory adaptations as increasing body size (Bergmann’s rule) and decreasing extremity length (Allen’s rule) helps minimize heat loss in cold environments (Mayr 1956; but see Scholander 1955). All skeletal phenotypes that were divergent between SARA and MANA were in accordance with Allen’s and Bergmann’s rules, with MANA mice having elongated limb bones and skull phenotypes, relative to their body size (Figure 1C-F, Supplementary File 3). The elongated rostrum of MANA mice may have functional importance in thermoregulation, as having a longer nasal cavity can lead to increased evaporative cooling as a method of heat dissipation (Walker and Wells 1961). This strict adherence to Allen’s and Bergmann’s rules suggests that adaptation to differing climates influenced morphological evolution between SARA and MANA mice across the entire skeleton.

In addition to length measurements, we investigated variation in skeletal shape and found that MANA mice have proportionally elongated distal limb elements (Supplementary Figure 1). Changes in limb proportions could be indicative of locomotor variation, with elongation of the distal limb associated with increased running endurance (Polly 2007). On the other hand, distal limb elements may play an outsized role in thermoregulation as they have a larger surface area to volume ratio than proximal elements, and thus modulating their length can help conserve or dissipate heat more effectively (Tilkens et al. 2007). Lastly, this pattern may reflect developmental differences, with distal elements being more evolutionary labile as a function of developmental timing (Stepanova and Womack 2020; Rothier et al. 2023). Functional studies would be needed to test these hypotheses.

### Genetic architecture of skeletal evolution is largely polygenic, with mostly additive QTL

The distribution of effect sizes for skeletal trait QTL included a few large-effect loci and many small-effect loci (Figure 3A). This genetic architecture fits an exponential distribution, matching the theoretical prediction of an adaptive walk toward a distant optimum with initial steps towards an optimum being mediated by large-effect loci, and subsequent steps mediated by many smaller-effect loci (Orr, 1998). However, the largest effect QTL identified only explained 4.7% of phenotypic variation and the average percent variation explained (PVE) was 1.7%. The largest PVE QTL identified here are less than that found in similar QTL studies (Miller et al. 2014) and below what is generally considered a “large-effect” QTL (20% PVE, Griswold, 2006). Ultimately, we found that skeletal evolution in tropical and temperate house mice is highly polygenic, with predominantly small-effect loci underlying phenotypic variation.

Second, we found that skeletal trait QTL had largely additive effects on phenotypic variation (Figure 3C). Newly arising beneficial mutations that are additive or dominant have a higher probability of fixation than recessive mutations and are thus expected to contribute more to adaptation (i.e. “Haldane’s sieve”; Haldane 1927). However, when selection is acting on standing genetic variation that was formerly in mutation-selection balance, Orr and Betancourt (2001) showed that fixation probability is independent of dominance. Previous analyses of introduced house mice in North America suggest that environmental adaptation in this system is largely the result of selection on preexisting genetic variation (Phifer-Rixey et al. 2018). Moreover, these studies have shown that adaptation seems to be due mainly to changes in gene regulation, and that much of this is due to *cis*-regulatory changes (Phifer-Rixey et al. 2018; Mack et al. 2018; Ballinger et al. 2023; Durkin et al. 2024). In general, *cis-*regulatory mutations are thought to be a primary driver of morphological adaptation (Carroll 2008; Wagner, Pavlicev, and Cheverud 2007; Stern 2000) and act additively (Lemos et al. 2008; McManus et al. 2010). Our findings of primarily additive loci underlying skeletal variation support this idea, and are similar to the dominance effect distributions reported in other examples of natural skeletal variation (Miller et al. 2014; Albert et al. 2008; Parmenter et al. 2016).

### Skeletal evolution is mediated by both modular, regionally specific loci and by several highly pleiotropic loci

The interconnected nature of the skeleton, and the oppositional forces of Allen’s and Bergmann’s rules, allowed us to address the extent of modularity and pleiotropy in trait evolution. Most QTL influenced only one or two traits, suggesting a genetic architecture of regional specificity with overall low levels of pleiotropy. Anatomical modularity is predicted to evolve during morphological divergence as a mechanism to avoid the negative pleiotropic effects of causing broad scale changes (Stern 2000; Carroll 2008). These changes could be controlled by modulating developmental genes that only operate on particular regions or through context-specific *cis*-regulatory changes. Similar studies in both house mice (Wagner et al. 2008) and stickleback fish (Albert et al. 2008; Miller et al. 2014) report similar levels of low pleiotropy and regionally specific changes.

However, we did identify several highly pleiotropic loci that influence traits across the skeleton. A quarter of all QTL identified are encompassed by these pleiotropic loci (influencing four or more traits). hpQTL on chromosomes 1 and 2 influence 10 of the 12 skeletal traits we analyzed (Figure 4). These QTL also mapped to PC1 (Figure 5B), which captures broad scale skeletal variation. An interesting aspect of these pleiotropic QTL was their unidirectional influence on phenotypic change (i.e. having the SARA allele for the hpQTL on chromosome 1 increased measurements for each trait it influenced, Figure 4D). Importantly, these effects were often opposite to the evolved change between MANA and SARA. Indeed, about half of all QTL showed antagonistic effects (Figure 2), which is higher than that observed for similar skeletal QTL studies in stickleback fish (Albert et al. 2008; Miller et al. 2014) and house mice (Parmenter et al. 2016). The high proportion of antagonistic QTL could be a signature of the concurrent, oppositional forces reflected in Allen’s and Bergmann’s rules and the compensatory changes needed to reach the unique fitness optima of each population. However, it is important to note that antagonistic effects could be the result of compensatory changes that “fine-tune” a phenotype via stabilizing selection after it overshoots the phenotypic optimum (Griswold & Whitlock, 2003).

There is growing evidence that loci of intermediate pleiotropy may be favored during adaptation as they could help facilitate the larger steps to a new fitness optimum required when organisms are facing novel selective pressures (Hämälä et al. 2020; Rennison and Peichel 2022). This may be particularly true in cases where selection is acting on standing genetic variation, where segregating alleles have already been subjected to selection (Thompson, Osmond, and Schluter 2019). However, the antagonistic effects of the hpQTL identified here are inconsistent with the idea that such highly pleotropic loci may facilitate adaptation. Instead, they may act as a constraining force. Perhaps this is particularly true in cases where discordant morphological changes are evolving concurrently. This is not to say that hpQTL do not confer a fitness advantage. In this scenario, hpQTL could be driven to high frequency via positive selection but for each population to reach its fitness optimum, additional compensatory changes would then also be necessary. It is important to note that we cannot distinguish truly pleiotropic loci from instances of tight genetic linkage given the low mapping resolution of QTL intercrosses. However, the nature of the hpQTL influencing phenotypes in the same direction suggests that they may represent one causal locus. Ultimately, while we identified several highly pleiotropic loci, most QTL only influenced one or two traits, providing evidence for a mix of broad scale pleiotropic and trait-specific changes in conferring adaptive phenotypes in this system.

### Identifying candidate genes underlying complex phenotypes

Undeniably, a primary goal of QTL mapping is to identify the causative genes underlying phenotypes of interest. However, the large size of QTL intervals precludes our ability to pinpoint causative genes. Indeed, the average number of genes in a single QTL interval in our study was 996 (651 for peaks identified only through single-QTL mapping, Supplementary File 5). There are several approaches to narrowing down such large lists to candidate genes of interest. First, QTL regions can be overlapped with genes that show signatures of selection in wild population, as we have done here. However, the overlap between our selection scan outliers and genes within PC QTL intervals was 343, a large enough list that finding promising candidates is still not feasible. Second, if there is a relevant tissue connected to the phenotype of interest, QTL regions can be narrowed down to only genes that are differentially expressed. Further, patterns of allele-specific expression in F1 individuals can identify expression differences that are controlled in *cis* and therefore linked to a causal locus controlling expression variation. Lastly, potential candidates can be identified through searching for genes with known phenotypic consequences in model systems. A great advantage of working with house mice is the rich database of known mutants (Baldarelli et al. 2024). Future work on gene expression in the developing skeletons of SARA and MANA mice at different developmental time points will be useful for identifying causative genes within the QTL intervals identified in the mapping population presented here.

## Data Availability Statement

The sequencing data in this article (fastq files for ddRAD-seq libraries) are available on NCBI under <u>BioProject PRJNA1295569</u>. Custom code used is available on GitHub (<u>s-durkin/Durkin_skeletalQTL_2025</u>). Phenotypic data and information on samples used are available in the supplementary files associated with this article.

## Acknowledgements

The authors are grateful for the help of many scientists: Craig Miller for advice on quantitative genetic analyses and helpful discussion; David Begun for providing comments on this manuscript; Alonso Corral, Marcella Welter, and Christina Gao for invaluable help in data collection; Lydia Smith and the Evolutionary Genomics Lab for guidance during ddRAD-seq library preparation; and Isaac Linn for guidance on statistical analyses, particularly relating to clustering of hpQTL.

## Figure and File Captions

**Supplementary Figure 1. Divergence in allometry indices in MANA and SARA mice**. A). Intermembral index (ratio of the forelimb to hindlimb [(HL + UL)/(FL + TL) x 100]). B). Humerofemoral index (ratio of proximal elements of the limb [(HL/FL) x 100]). C). Crural index (ratio of distal to proximal elements of the hindlimb [(TL/FL) x 100]). D). Brachial index (ratio of distal to proximal elements of the forelimb [(UL/HL) x 100]). P-values represent Welch two sample t-tests.

**Supplementary Figure 2. Distribution of QTL effect sizes for individual traits.** For each plot, the X-axis is percent variation explained, and the Y-axis corresponds to the chromosome of each QTL.

**Supplementary Figure 3. Pairwise overlap enrichment of QTL identified in single-QTL mapping.** Simulated number of pairwise overlaps per chromosome. Red dot represents the observed value of pairwise overlaps. Chromosomes that contained more overlaps than expected by chance (observed value beyond the top 5% of simulated distribution) are colored purple. Only peaks identified in the single-QTL, scanone mapping approach are included.

**Supplementary Figure 4. Mode of action for hpQTL on chromosomes 7 and 9**. Left y-axis and purple bars display the effect size and direction of the hpQTL on each trait it influences; negative values correspond to the SARA allele conferring a higher phenotype value, and vice versa for the MANA allele. Right y-axis and black points display the phenotypic difference between MANA and SARA, for residual phenotype values.

**Supplementary Figure 5. Investigation of recombination rate in QTL intervals.** A). Distribution of recombination rate for each chromosome estimated in 5Mb windows across the genome. Red dot represents the average recombination rate within QTL intervals on that chromosome. No QTL intervals had an elevated or depressed recombination rate as compared to the background average. B). Relationship between effect size and recombination rate of individual QTL. Correlation coefficient and p-value are Pearson’s R.

**Supplementary Figure 6. QTL intervals for principal components without body weight as a covariate.** Significant QTL intervals for PC1 (A) and PC2 (B). Dark purple QTL were identified through the single-QTL mapping approach, while light purple intervals correspond to QTL identified only through the multi-QTL mapping approach. Body weight was not included as a covariate.

**Supplementary Figure 7. Significant overlap between PBSn1 outliers and genes in PC QTL intervals**. Expected number vs. simulated distribution of overlap between genes within PC QTL intervals and PBSn1 outliers in either the NH/VT or Manaus test at the 1% (A), 5% (B) and 10% (C) outlier level. Dashed line indicates the observed overlap of PC and PBSn1 outlier genes. All overlaps are more than expected by chance (observed value beyond the top 5% of simulated distribution).

**Supplementary File 1.** Phenotype measurements for all skeletal traits and body weight for F3 and parent mice.

**Supplementary File 2.** Details on skeletal measurements and landmarks used.

**Supplementary File 3.** Mean phenotype values and Student’s t-test results for parental trait divergence.

**Supplementary File 4.** Location, QTL id, effect size, dominance, and detection method information for all QTL peaks.

**Supplementary File 5.** Number of genes in the GRCm39 mouse genome within QTL regions for all identified QTL.

